# Optical inhibition of zebrafish behavior with anion channelrhodopsins

**DOI:** 10.1101/158899

**Authors:** Gadisti Aisha Mohamed, Ruey-Kuang Cheng, Joses Ho, Seetha Krishnan, Farhan Mohammad, Adam Claridge-Chang, Suresh Jesuthasan

## Abstract

In behavioral analysis, optical electrical silencing provides a way to test neuronal necessity. Two light-gated anion channels, GtACR1 and GtACR2, have recently been shown—in neuronal culture and in *Drosophila*—to inhibit neurons potently. Here, we test the usefulness of these channels in zebrafish. When the GtACRs were expressed in motor neurons and actuated with blue or green light, fish spontaneous movement was inhibited. In GtACR1-expressing fish, only 3 µW/mm^2^ of light was sufficient to have an effect; GtACR2, which is poorly trafficked, required stronger illumination. After light offset, GtACR-expressing fish movement increased; this suggested that termination of light-induced neural inhibition may lead to depolarization. Consistent with this, two-photon imaging of spinal neurons showed that intracellular calcium also increased following light offset. The activity elicited at light offset needs to be taken into consideration in experimental design, although this property may help provide insight into the effects of stimulating a circuit transiently. These results show that GtACR1 and GtACR2 can be used to optically inhibit neurons in zebrafish, and thus to test neural circuit necessity.

## Introduction

One approach to understanding the role of specific neurons in a given behavior is to experimentally alter their activity. A precise means of doing this is by using light-gated channels [1–4], Optogenetic activators such as Channelrhodopsin-2 [5], ChlEF [6] and CsChrimson [7] can reversibly depolarize membrane potential with millisecond resolution, and have been widely used in a range of organisms [8–11]. For numerous different behavioral functions, these tools have enabled the determination of neuronal sufficiency. While optical activation can demonstrate sufficiency, establishing whether neurons are normally involved requires additional experiments. Optical or electrical recording can determine which activity patterns are correlated with behavior. However, for the crucially important loss of function experiment, determining necessity requires effective tools for neuronal inhibition.

A number of different optogenetic inhibitors have been developed. The first generation of these tools include light-actuated pumps like halorhodopsin [12,13], which facilitates light-dependent chloride entry, and archaerhodopsin [14,15], a proton pump that is also hyperpolarizing. A limitation of these molecules is their low conductance and the high levels of expression and illumination that are required for effective silencing. For example, the archaerhodopsin derivative Archerl requires 3 mW/mm^2^ of light to inhibit action potentials [16]. A second generation of optogenetic inhibitors includes genetically modified channelrhodopsins such as ChloC [17] and iC1C2 [18], which hyperpolarize neurons by light-gated conductance of chloride. Both require similar levels of illumination, although an improved version of ChloC, iChloC, requires 10 times less light [19]. More recently, naturally evolved anion-conducting channels from the alga *Guillardia theta* were shown to silence neurons at very low light levels, i.e. in the range of µW/mm^2^ [20]. *Drosophila* experiments have confirmed that these optogenetic tools are potent inhibitors *in vivo* [21]. Here we extend these results to larval zebrafish, to establish that *Guillardia theta* anion channelrhodopsins are effective inhibitors of neural activity in a genetically-tractable vertebrate.

## Materials and Methods

### Generation of GtACRl and GtACR2 transgenic zebrafish lines

Sequences encoding GtACRl and GtACR2 fused to eYFP [21] were placed downstream of the upstream activating sequence (UAS) using Gateway cloning (Thermo Fisher Scientific). The resulting construct was cloned into a plasmid containing Tol2 sequences to facilitate integration into the zebrafish genome [22,23]. The constructs (33 ng/ul) were then injected into nacre^-/-^ eggs at the one cell stage, along with elavl3:Gal4 DNA (15 ng/ul) to induce GtACR expression, and Tol2 mRNA (33 ng/ ul) to facilitate genomic integration. Embryos were screened after 24-36 hours for eYFP expression. Healthy embryos with eYFP expression grown to adulthood. At two months, fish were fin-clipped and PCR-screened with GtACR specific primers (GtACRl Forward: 5′-CACCGTGTTCGGCATCAC-3′, GtACRl Reverse: 5′-G CC ACC ACC AT CT CG A AG-3′, GtACR2 Forward: 5′-AT-TACCGCTACCATCTCCCC-3′, GtACR2 Reverse: 5′-TGGT-GAACACCACGCTGTAT-3′) to test for the presence of transgene. This led to the generation of two transgenic lines, *Tg(UAS:ACRl-eYFP)sq211* and *Tg(UAS:ACR2:eYFP)sq212.*

### Confocal imaging

24-hour old F1 embryos were dechorionated, anaesthetised with 160 mg/L tricaine, and mounted in 1% low melting agarose in E3. Imaging was carried out using a Zeiss LSM800 confocal microscope with a 40x water immersion objective.

### Spontaneous movement and light stimulation

*Tg(sl020t:GAL4, UAS:ACR1-eYFP)* and *Tg(sl020t:GAL4, UAS:ACR2-eYFP)* fish were screened with a fluorescence stere^-^ omicroscope at 23-24 hours post fertilization to identify ACR-expressing fish eYFP in the spinal cord. Embryos, still within their chorions, were then placed in a glass dish with 24 concave wells on a stereomicroscope (Zeiss Stemi 2000) with a transmitted light base. Behavior was recorded on the microscope using a Point Gray Flea2 camera controlled by MicroManager. Stimulating light was delivered by LED backlights (TMS Lite), with peak intensity at 470 nm (blue), 525 nm (green), or 630nm (Red), placed adjacent to the glass dish. 595 nm (amber) illumination was provided using LEDs from CREE (XR7090-AM-L1-0001), which were mounted onto thermal LED holders (803122; Bergquist Company). The intensity of light was measured using a S120VC power sensor and a PM100A console (Thorlabs). LEDs were switched on and off using an Arduino board controlled by MicroManager, to regulate the power supply unit. Embryos were recorded for a total of 45 seconds, with the LED being turned on 15 seconds after the start of recording and turned off at 15 seconds later.

### Analysis of behaviour recordings

Image analysis was carried out using FIJI (RRID:SCR_002285) [24] as well as scripts written in Python. From the raw recordings, one frame was extracted per second to obtain a total of 46 frames (including the first and last frame). Circular regions of interest (ROIs) were manually drawn around each chorion to isolate each fish. Each frame was then subtracted from the next frame to identify the differences between frames. After thresholding, the number of different pixels in each ROI was taken as a measure of movement of each embryo [21]. An embryo was considered to have active movement, rather than to drift within the chorion, if there were more than 15 pixels that differed between frames. Any embryo that did not move during the entire recording was discarded from analysis. Estimation statistical methods were employed to analyze mean differences between control and experimental groups [25–27]. 95% confidence intervals for the mean difference were calculated using bootstrap methods [28]. All confidence intervals were bias-corrected and accelerated [29], with resampling performed 5,000 times. All reported *P* values are the results of Wilcoxon *t*-tests.

### Two-photon calcium imaging

Triple transgenic zebrafish embryos (s1020t:GAL4, UAS:ACR1-eYFP, elavl3:GCaMP6f) were mounted in agarose (2% low melting temperature, in E3). To enable individual cells to be followed without motion artifact, fish were anesthetized with mivacurium chloride (Mivacron, GSK). All imaging was performed using an upright Nikon A1RMP two-photon microscope equipped with a 25x 1.1 NA water immersion objective. Images were captured at a rate of 1 Hz, with the laser tuned to 920 nm. Blue light was delivered with the same light box used for behavior experiments, at the maximum intensity. Green light was not used, as this would overlap with the emission wavelength of GCaMP6f.

### Analysis of calcium-imaging data

Analysis was carried out using FIJI [24], unless otherwise stated. Background correction was first performed by subtracting the average value of a region outside the embryo for each frame. This was to eliminate the bleed-through from the illuminating LED. A median filter with radius of 1 pixel was then applied. Images were registered using MetaMorph. Regions of interest were drawn manually around cells. Average fluorescence intensity of cells within the ROI was obtained by measuring only pixels above a threshold.

## Results

### Transgenic zebrafish express GtACRl and GtACR2 in neurons

To express the anion channelrhodopsins in zebrafish, transgenic lines containing the coding sequences under the control of the upstream activating sequence (UAS) were generated using Tol2-mediated transgenesis. When crossed with GAL4 drivers, expression of the protein could be detected by the eYFP tag (Figure 1a, b). GtACRl-eYFP was detected in the periphery of the cell (Figure 1c), consistent with membrane localization. Peripheral labelling with GtACR2-eYFP could not be seen, although puncta could be detected (Fig. 1d), suggesting that GtACR2-eYFP was aggregating or was not efficiently trafficked to the plasma membrane.

**Figure 1.**
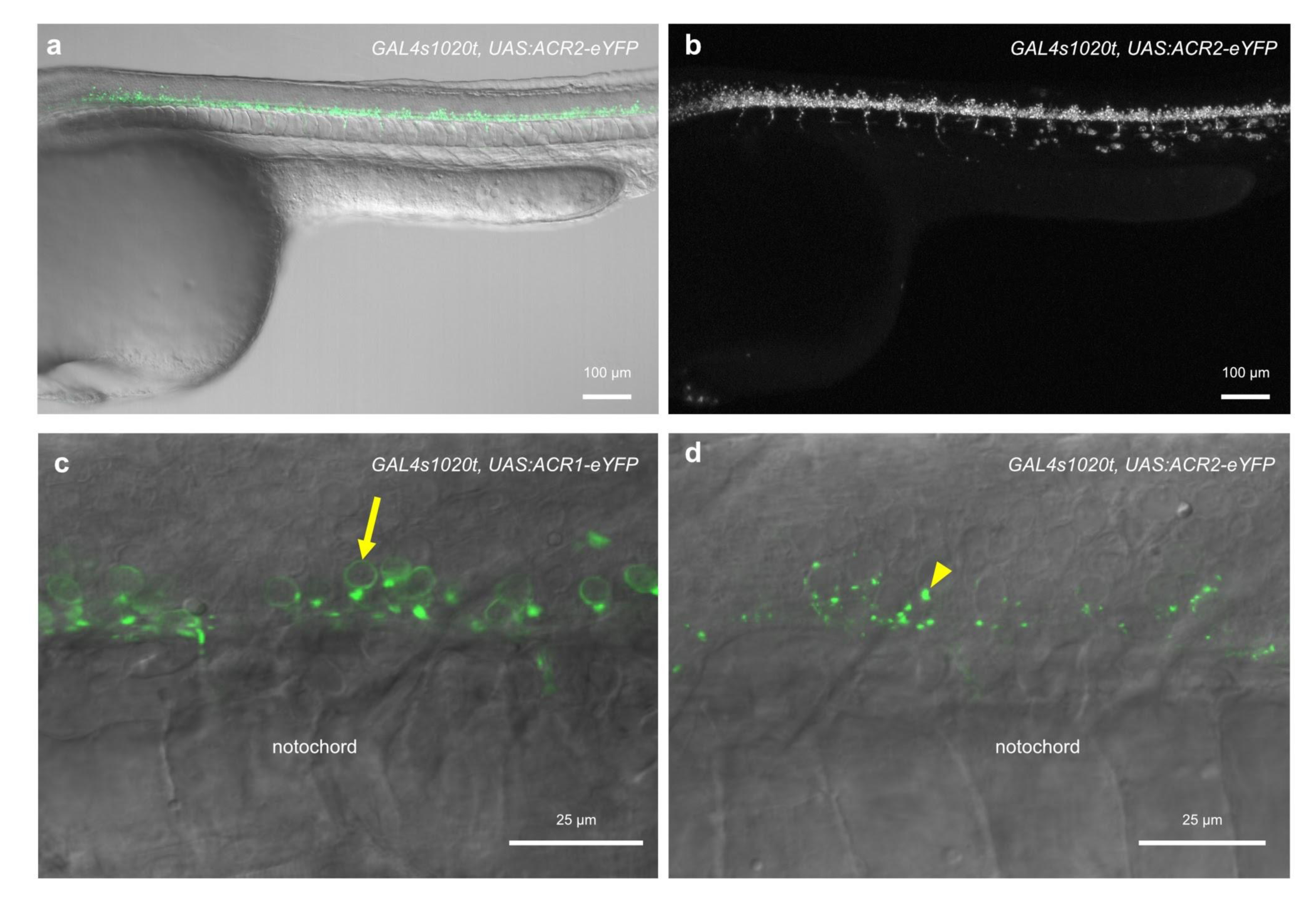
Expression of ACR1 and ACR2 in transgenic zebrafish. a, b. Expression of ACR2-eYFP in the trunk of a ~30 hpf *sl020t:GAL4, UAS:ACR2-eYFP* embryo, **c, d** High magnification view of the trunk of fish expressing ACRl-eYFP **(c)** or ACR2-eYFP **(d)** in spinal neurons. There is label at the cell periphery (arrow), indicative of localization to the plasma membrane for ACR1. In the case of ACR2, puncta are visible (arrowhead). Gamma = 0.45 for panel **b.** Anterior is to the left. Panels **a, c** and **d** are single planes, while **b** is a maximum projection.

### Light-actuated GtACRl and GtACR2 inhibit spontaneous movement

Between 17 and 27 hours post-fertilization, zebrafish display spontaneous coiling movements [30] (Figure 2a, b) that are dependent on the activity of motor neurons [31]. To test the ability of GtACRl and GtACR2 to inhibit coiling behavior, we measured the effect of light on the spontaneous movements of fish expressing these channels in motor neurons. Fish were exposed to a 15 s pulse of light in the middle of a recording lasting 45 s, and the number of fish moving in each 1 s bin was counted. With both ACR1 and ACR2, green (10 µW/mm^2^) and blue (14 µW/mm^2^) light caused a visible reduction in the proportion of animals showing spontaneous movement (Fig. 2c,d). Red light (9 µW/mm^2^) did not have a clear effect. This suggests that anion channelrhodopsins could be used for behavioral inhibition in larval zebrafish, with blue and green light.

**Figure 2.**
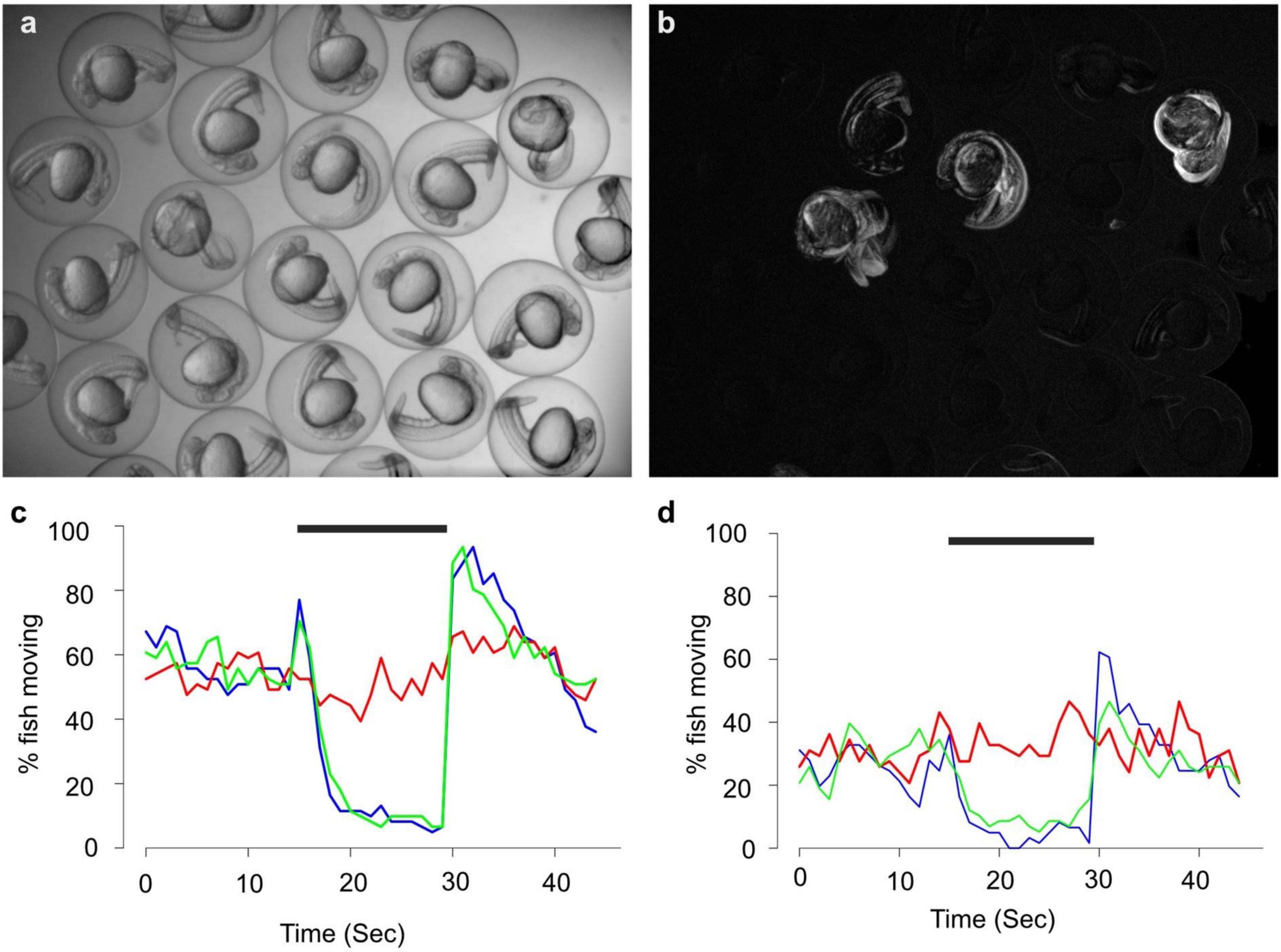
Spontaneous coiling movements as an assay for optical inhibition. a. 25 hour old zebrafish embryos, within their chorions. **b** Spontaneous movement over 10 seconds, as seen by summing together the difference between 10 consecutive frames. **c,d** Percentage of GtACRl **(c)** and GtACR2 **(d)** expressing embryos exhibiting spontaneous movement in each second, over a 45 second period. Fish were illuminated with blue, green or red light in the middle of the recording, in the region indicated by the black bar. The color of the line indicates the illumination wavelength. The intensities used are ~ 10 µW/mm^2^(blue), 14 µW/mm^2^ (green) and 9 µW/mm^2^ (red).

We next tested the effect of different intensities of light (Figure 3), to obtain a measure of the sensitivity of the transgenic lines to optical control. Here, the amount of movement per animal was measured in terms of pixel value differences between subsequent frames. High and medium intensities of green ( and blue light had a similar effect on ACR1-expressing fish (Figs. 3a,b, 4a; Table 1). Low intensities (3.3 µW/mm^2^ for blue and 2.3 µW/mm^2^ for green) were able to cause freezing, but at reduced efficiency compared to the higher intensities. For ACR2-expressing fish, high and medium intensities of blue light were able to induce similar levels of freezing in the zebrafish larvae, but only green light at high intensity was able to inhibit larval movement (Fig. 3c, d; Table 2). This suggests that ACR1 allows more effective behavioral inhibition.

**Figure 3.**
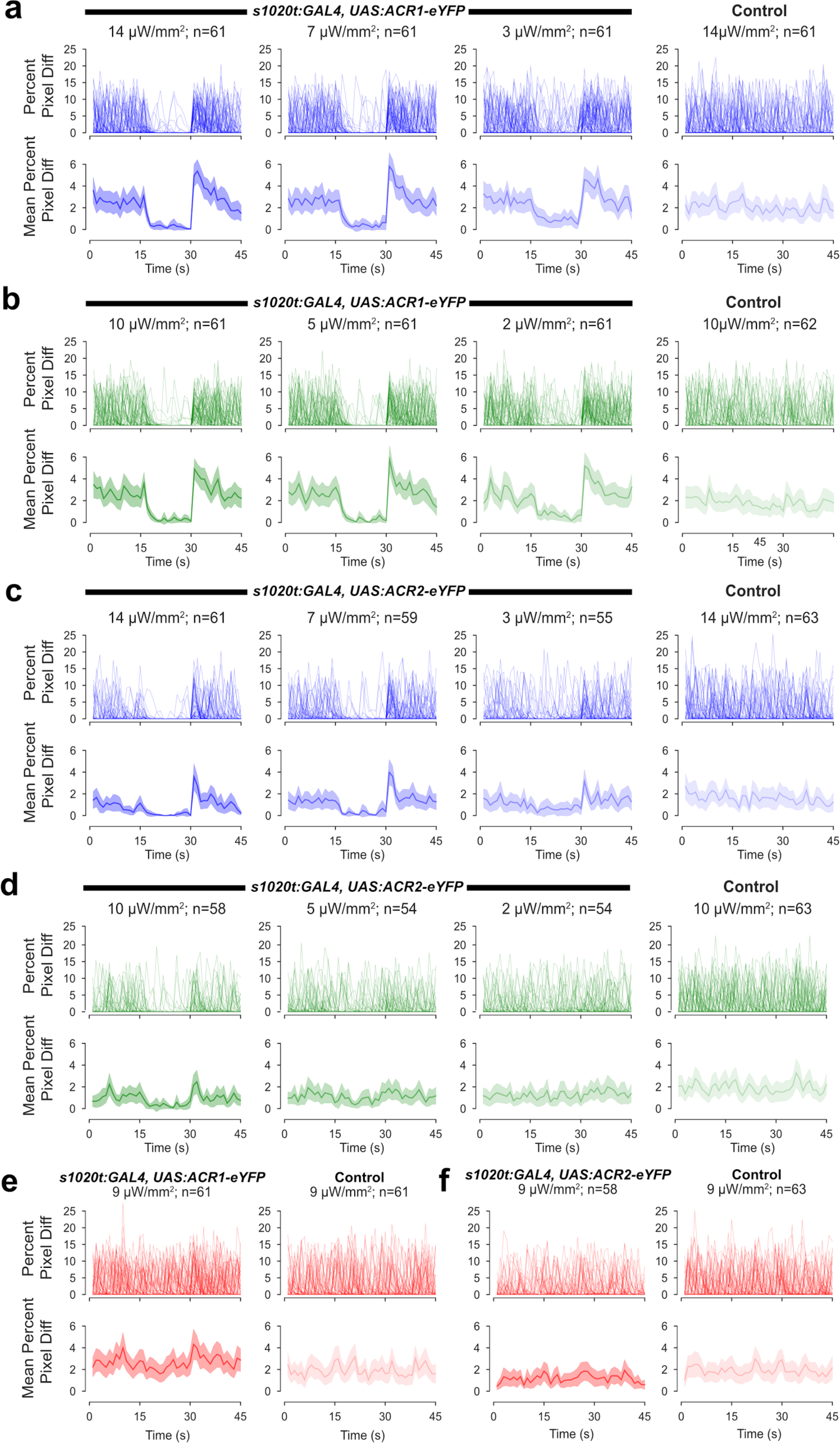
Light sensitivity of ACR1 and ACR2 expressing fish. a, b. Response of ACR1 fish to blue **(a)** and green **(b)** light of three different intensities (high, medium and low). For each panel, the top trace shows the movement of each fish in 1 second. This is measured by number of pixels that differ, relative to the cross-sectional area of the chorion. Each line represents one fish. The lower trace shows the mean value and 95% confidence intervals. Controls refer to siblings lacking eYFP expression that were exposed to high intensity light. **c, d** Effect of blue **(c)** and green **(d)** light of different intensities on ACR2 expressing fish. **e, f** The effect of red light on ACR1 **(e)** and ACR2 **(f)** expressing fish. Line color corresponds to the color of light used for illumination. For all conditions, the period of illumination lasted from t=15s to t=30s.

**Figure 4.**
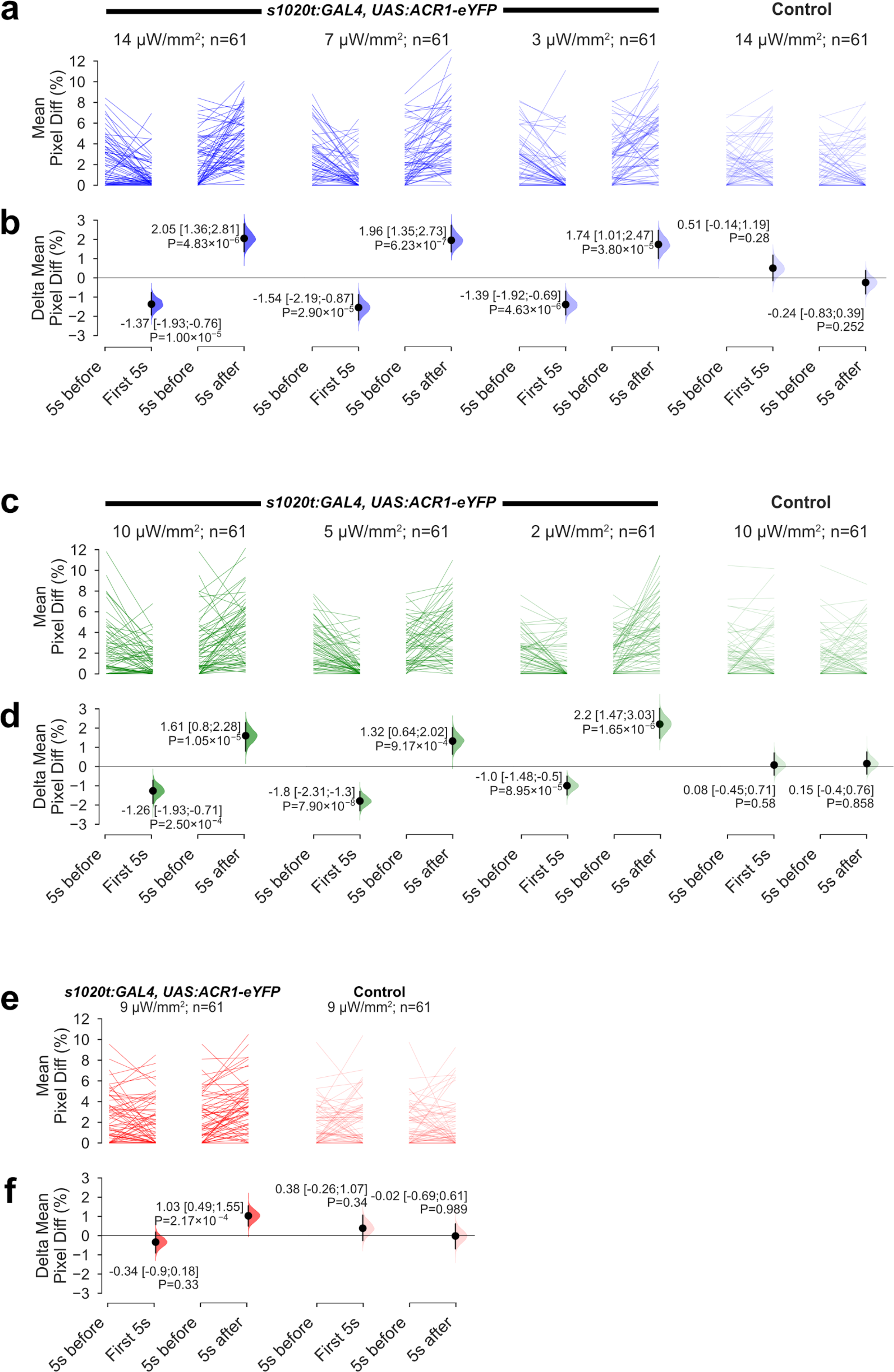
The effect of light and loss of light on spontaneous movement of GtACRl embryos. **a, c, e.** The amount of movement displayed by individual embryos, in the 5 seconds before light onset (–5s before–), in the first five seconds after light onset (–First 5s–), and in the first five seconds after light offset (–5s after–). Different intensities of blue **(a),** green **(c)** and red **(e)** light were tested. Controls refer to siblings lacking eYFP expression. The line color of each plot corresponds to the light color employed for illumination, **b, d, f.** The mean amount of movement, relative to the period before light onset. A negative value indicates inhibition of spontaneous movement, whereas a positive value indicates elevated levels of movement. Mean differences and 95% CIs are reported alongside *P* values from Wilcoxon tests. Blue **(b)** and green **(d)** light both affect spontaneous movement. Red light has a small effect **(f).**

For GtACRl, even the lowest intensity of blue (3.3 µW/mm^2^) or green (2.3 µW/mm^2^) light caused a reduction in movement (Fig. 4a, b). With GtACR2, behavioral inhibition was seen with low intensity of blue light (Fig. 5a, b), but not with green light (Fig. 5c, d). Red light did not inhibit coiling of either GtACRl or GtACR2 fish at the intensity tested (Fig. 3e, 4e), consistent with the reported action spectrum of both GtACRl and GfACR2 (Ref).

**Figure 5.**
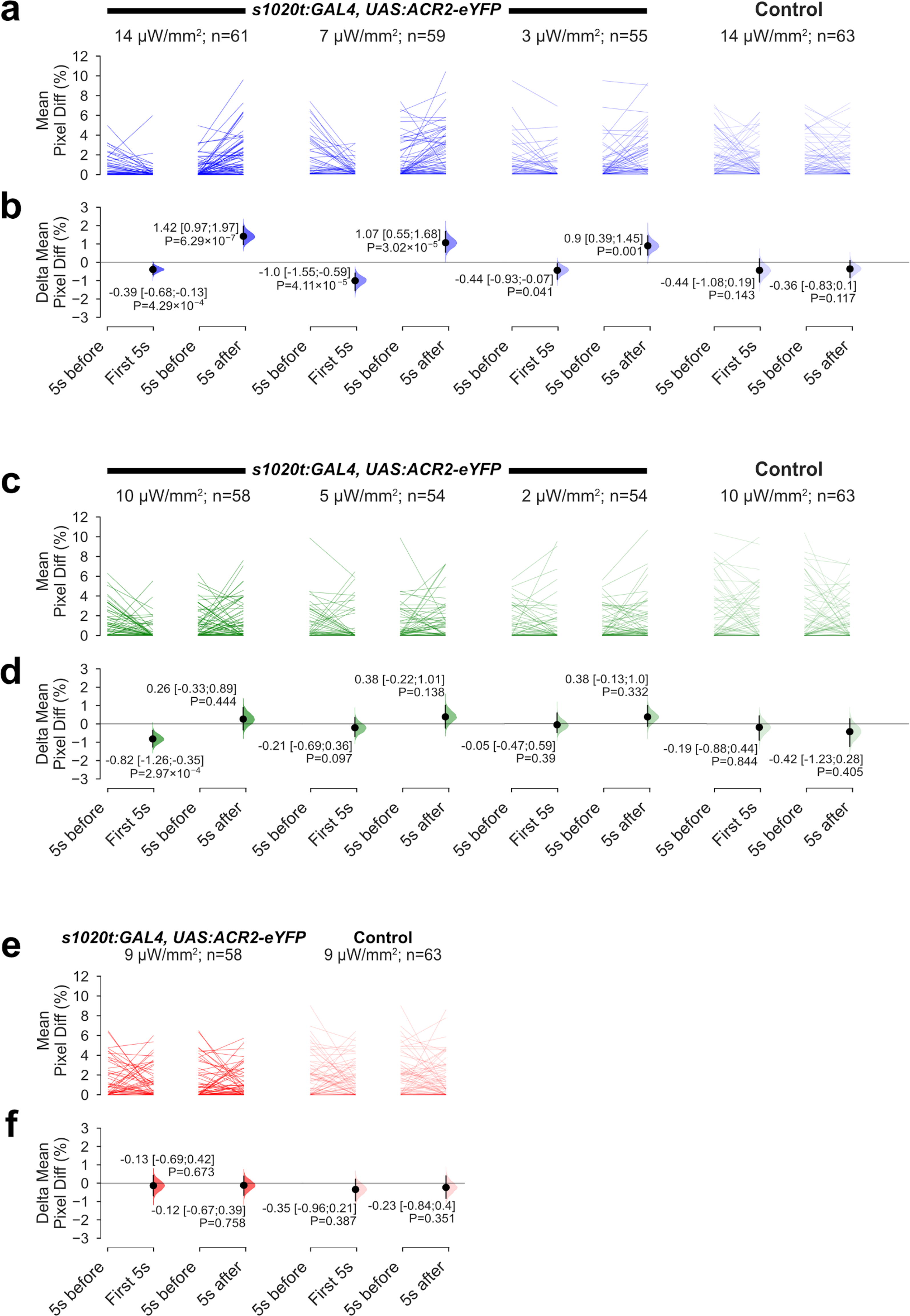
The effect of light and loss of light on spontaneous movement of GtACR2 embryos. a, c, e. The amount of movement displayed by individual embryos, in the 5 seconds before light onset (–5s before–), in the first five seconds after light onset (–First 5s–), and in the first five seconds after light offset (–5s after–). Three different intensities of blue **(a)** and green **(c)** light, and one intensity of red light **(e)** were tested. The color of the lines indicates the color of light used for illumination. Controls for each experiment were siblings not expressing eYFP. **b, d, f.** The mean amount of movement, relative to the period before light onset. A negative value indicates inhibition of spontaneous movement, whereas a positive value indicates elevated levels of movement. Mean differences and 95% CIs are reported; all *P* values are results of Wilcoxon tests. Blue **(b)** light affected spontaneous movement at medium and high intensities, whereas green light only had a weak effect had high intensity **(d).** Red light had no effect.

### GtACRs increase behavioral activity at light offset

When the actuating light was turned off, a large percentage of GtACRI and GtACR2-expressing embryos displayed prolonged coiling movement (Fig. 2c,d). To assess whether this reflected an increased amount of movement per animal, or an increased probability but similar level of movement, we looked at the response of each fish, comparing movement in the 5 s period after light offset with the 5 s period before light onset. As can be seen in Figures 4 and 5, there is a greater amount of movement following light offset, implying that loss of light triggers stronger movement than what is seen spontaneously.

This effect was observed in larvae expressing GtACRI in both green and blue light, at all three light intensities tested (Fig. 4). However, for larvae expressing GtACR2, this effect was only observed upon exposure to blue light and strong green light. This suggests that the movement to light offset is not due purely to change in illumination (i.e. a startle response), but is due to presence of the light gated anion channel.

Halorhodopsin has previously been used to inhibit neural activity in zebrafish larvae [32]. We compared the performance of this chloride pump with the anion channelrhodopsins. At the intensities that ACRs enabled a behavioral manipulation of zebrafish, no light-evoked inhibition of spontaneous movement, or induction of movement at light offset, was seen with halorhodopsin (Fig. 6). Thus, anion channel rhodopsins are more effective tools for light-gated control of behavior in zebrafish, compared with halorhodopsin.

**Figure 6.**
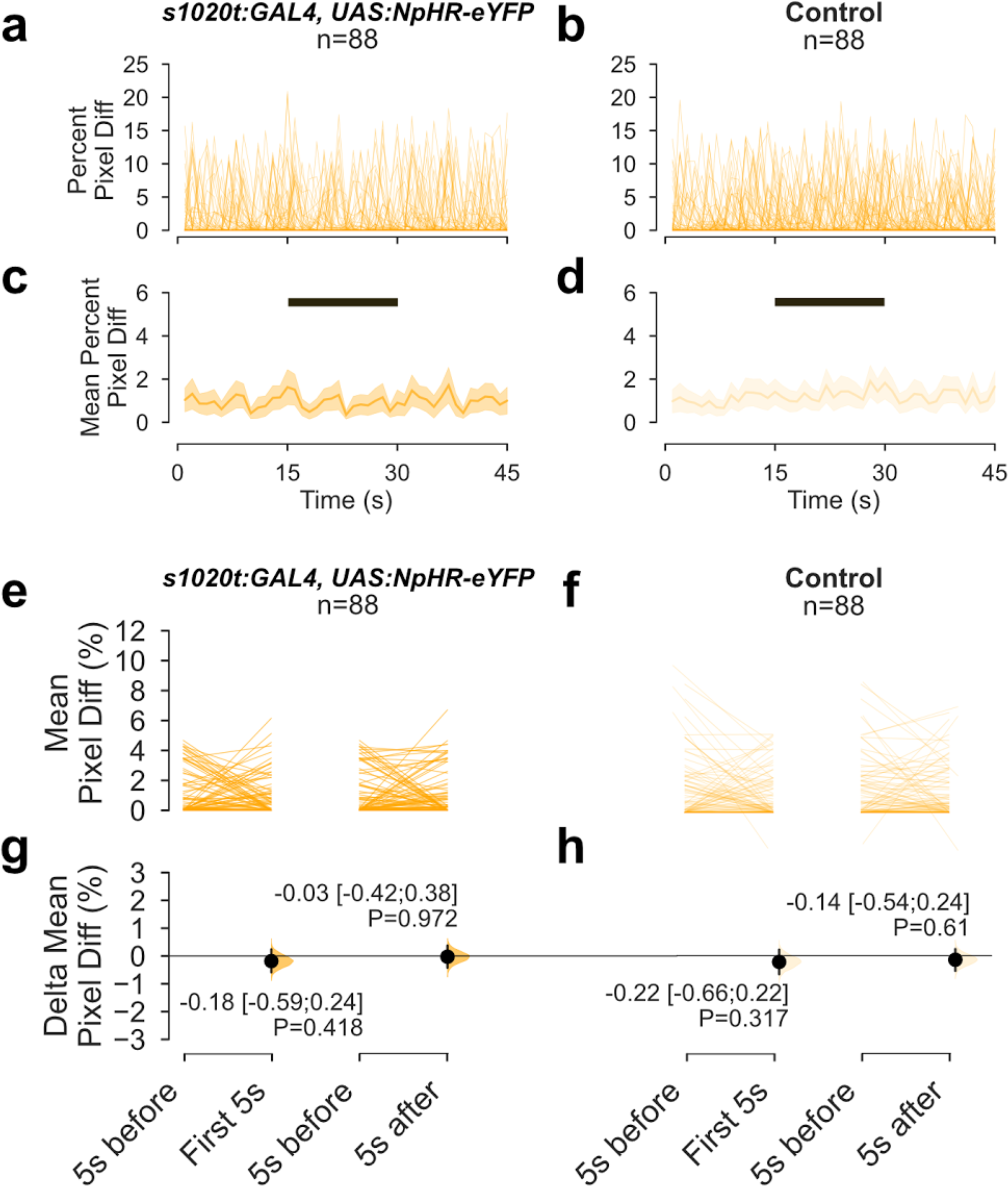
The effect of light on spontaneous movement of NpHR-expressing embryos, a-d. Time-course of movement of embryos, with **(a,c)** or without **(b,d)** halorhodopsin expression, in a 45 second recording with exposure to 15 seconds of amber light. The period of illumination is indicated by the black bar. **a** and **b** show movement of individual embryos, while **c** and **d** show mean values and 95% confidence intervals, **e, f.** Average amount of movement in the 5 seconds before light onset (–5s before–), in the first five seconds after light onset (–First 5s–), and in the first five seconds after light offset (–5s after–), in NpHR-expressing embryos **(e)** and non-expressing siblings **(f). g, h.** Difference in movement in the first five seconds after light onset and after light offset, relative to the period before light, in NpHR-expressing **(g)** and non-expressing **(h)** embryos. Intensity of amber light used = 17 µW/mm^2^.

### Calcium imaging of spinal neurons

As an independent method of assessing the effect of light and loss of light on neurons in zebrafish larvae expressing GtACRl, two-photon calcium imaging was carried out. We used the *elavl3* promoter to drive broad neuronal expression of the calcium indicator GCaMP6f. In non-anesthetized *Tg(elavl3:GCaMP6f, sl020t:-GAL4, UAS:ACR1)* fish, fluorescence levels in spinal neurons rose when spontaneous movements occurred (Fig. 7a, b; Movie 1). In the presence of blue light, however, there were no spikes of GCaMP6f fluorescence and movement ceased. At the offset of blue light, calcium levels and movement both increased. In siblings that did not express ACR1, increase in calcium levels persisted in the presence of light (Fig. 7c,d). These observations are consistent with previous observations that spontaneous movement is accompanied by a rise in intracellular calcium [33], and also suggest that offset of the actuating light may drive neural firing in fish expressing ACR1. To investigate this further, we used focused on fish at a later stage where spontaneous activity was not as prominent, so that signals evoked by loss of light can be clearly distinguished from spontaneous activity. As seen in Fig. 7e, f, spinal neurons that had no activity before light showed an increase in fluorescence after the offset of blue light. This observation suggests that the termination of light-gated silencing can lead to depolarization of neurons within the spinal network.

**Figure 7.**
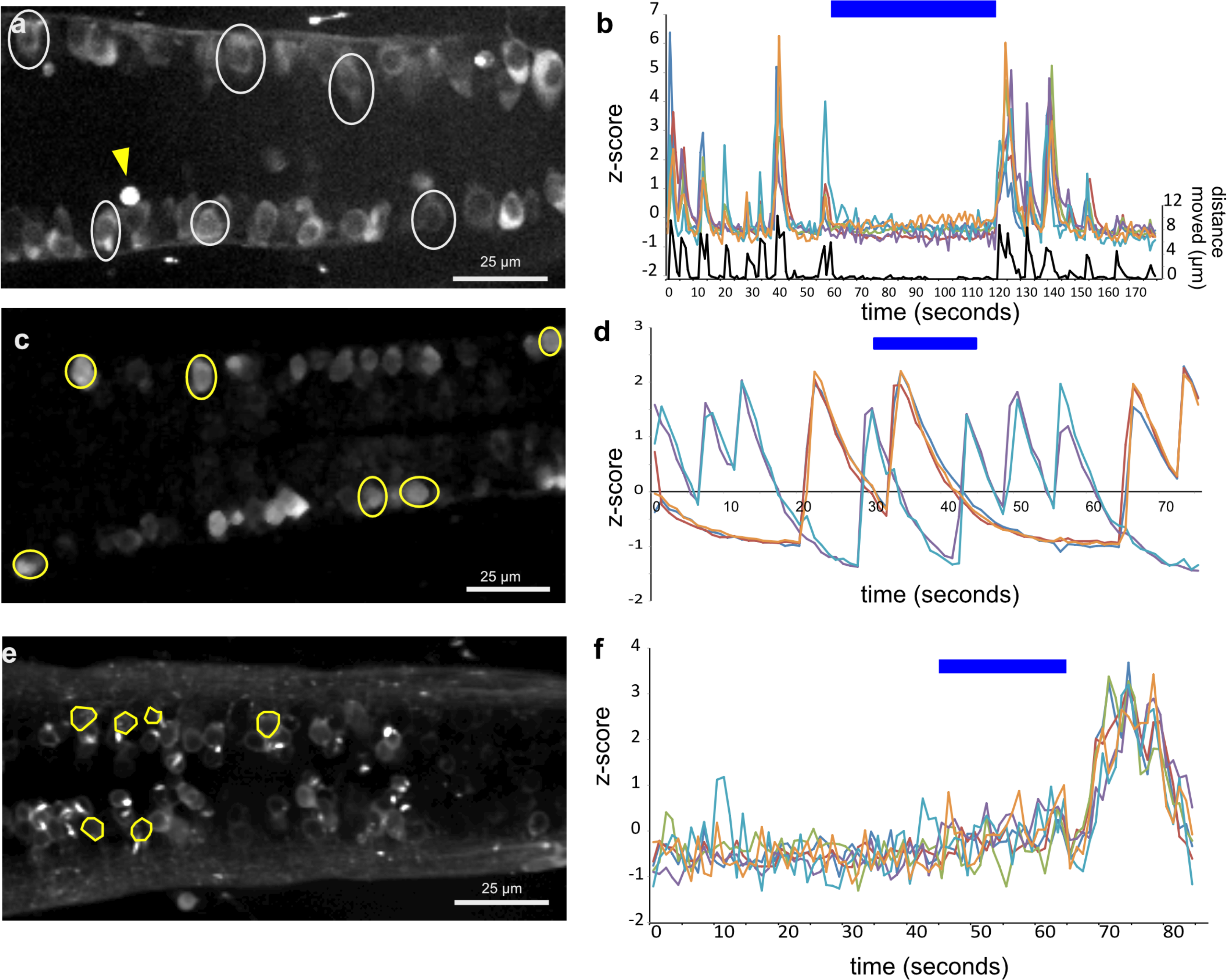
Calcium imaging of spinal neurons in ACR1 embryos and non-expressing siblings. a. Dorsal spinal neurons of a 24 hour old ACR1 embryo expressing GCaMP6f under the *elavB* promoter, **b.** Time-course of fluorescence intensity from neurons, represented as z-scores, and movement (magenta trace) as measured by tracking a bright structure (arrowhead in a). The period of blue light illumination is indicated by a bar at the top of the chart. **c,d.** The response of six neurons in a 24 hour non-ACRl embryo, outlined in yellow **(c)** to blue light, **d.** Increase in fluorescence intensity occurs during the period of illumination with blue light, **e, f** The response of five spinal neurons in a 4 day old ACR1 embryo. There is no activity before light in these neurons, but there is a rise in GCaMP6f fluorescence after termination of the blue light.

## Discussion

We have investigated the usefulness of anion channelrhodopsins as a tool for optical control of zebrafish behavior. By expressing these channels in spinal neurons of zebrafish and exposing the animals to light, spontaneous coiling movements could be completely and reversibly inhibited. GtACRl appears to be a more effective tool, as ACR1-expressing fish were affected by both green and blue light at the lowest intensity tested, which is ~3 µW/mm^2^. GtACR2 was able to inhibit movement, but was effective mainly with blue light at medium or high intensity; green light could inhibit movement only at high intensity. Halorhodopsin, which has been used previously in zebrafish [32], had no effect on spontaneous movement at low intensity tested here. This suggests that, as in *Drosophila [21]*, the anion channelrhodopsins are potent tools for the light-mediated inhibition of neural activity in zebrafish.

The poorer performance of GtACR2 compared to GtACRl may be due to this molecule’s poor membrane localization. As chloride channels, the function of the GtACRs requires that the proteins be localized to the plasma membrane. One approach that can be taken to improve the performance of GtACR2 could thus be to add a trafficking signal from Kir2.1, as was done with halorhodopsin to create eNpHR3.0 [34]. This may enable the advantages of GtACR2, i.e. it is a blue-shifted action spectrum and faster kinetics [20], to be used in the zebrafish system.

The termination of light-evoked silencing appears to lead to depolarization of neurons, as judged by increased coiling behavior as well as rise in intracellular calcium. This property of ACRs has been observed in other light-gated chloride pumps and is linked to an increase in excitability due to accumulation of chloride ions inside the cell which elevates mean spike probability and mean stimulus-evoked spike rate by changing the reversal potential of the GABAA receptor [35]. Thus, in designing experiments where the goal is to test the effects of silencing a particular set of neurons, it may be advisable to restrict observations to the period during which light is delivered. The burst of activity that occurs after the offset of light may be problematic in study of long-term processes, such as memory [36] or emotion [37]. For acute processes, however, this property may be beneficial as it provides a way to test the effects of activating the same set of neurons. An example of this is shown in the companion manuscript (Cheng *et* al.), where the direction of swimming is reversed in light versus darkness when ACRs are expressed in the thalamus.

## Conclusion

The anion channelrhodopsins GtACRl and GtACR2 provide useful tools for optical manipulation of neural circuits in zebrafish.

**Movie 1. The effects of blue light on the trunk of** *s1020t:GAL4, UAS:ACR1, elavl3:GCaMP6f* **fish.** Twitches of the body are accompanied by increase in GCaMP6f fluorescence. In the presence of blue light, which can be seen by an overall increase in brightness, the embryo no longer twitches. After the blue light is switched off, movement and change in GCaMP6f fluorescence resume. This is a dorsal view, with anterior to the left.

## Author contributions

*Conceptualization:* SJ and ACC; *Methodology:* GAM, SK, SJ; *Software:* JH (Python); *Investigation:* GAM (behavior, transgenic design, genetics), SJ (confocal microscopy) and CRK (calcium imaging), *Resources:* FM (synthetic GtACR genes); *Data Analysis:* GAM, FM and JH (behavior), and SJ (calcium imaging); *Writing – Original Draft:* SJ and GAM with contributions from all authors; *Writing – Revision:* SJ and ACC; *Visualization:* JH and SJ; *Supervision:* SJ and ACC; *Project Administration:* SJ and ACC; *Funding Acquisition: ACC* and SJ.

## Acknowledgements

We thank John Spudich (The University of Texas Medical School at Houston) for sharing the GtACR sequences. Major support for GAM, RKC, SK and SJ was from a start-up grant from the Singapore Ministry of Education to SJ. ACC and SJ were supported by A*STAR Joint Council Office grant 1431AFG120. JH was supported by the A*STAR Scientific Scholars Fund. FM and ACC received support from Duke-NUS Medical School and Ministry of Education grant MOE-2013-T2-2-054. SK was supported by an NUS Graduate School for Integrative Sciences and Engineering (NGS) Scholarship. The authors were supported by a Biomedical Research Council block grant to the Institute of Molecular and Cell Biology.

